# Shear Difference: Flow Type Dictates Endothelial Flow-Responsive Gene Programs in a 3D-Printed in vitro Model

**DOI:** 10.1101/2025.11.24.690338

**Authors:** NA Shah, KA Rye, ZH Endre, TJ Barber, JH Erlich, BJ Cochran

## Abstract

**Background:** Endothelial cells (ECs) are mechanosensitive and adopt distinct phenotypes in response to hemodynamic forces. These phenotypes are rarely seen in shear-responsive signalling studies because commonly used in vitro platforms rarely reproduce the true scale of vessel geometries or physiologic waveforms.

**Aim:** To determine how the presence and temporal pattern of flow impact endothelial morphology and transcriptional activity in a 3D macrofluidic model.

**Methods:** Idealised vessels were 3D-printed using a water-soluble polyvinyl alcohol filament and cast in polydimethylsiloxane. The polyvinyl alcohol cores were dissolved leaving a polydimethylsiloxane lumen on which HMEC-1 cells were grown and perfused for 24 h under static, continuous flow, or pulsatile flow. Cell morphology was assessed using immunofluorescence. Bulk RNA-seq was performed with Hallmark pathway enrichment using gene set enrichment analysis.

**Results:** Relative to static culture, continuous flow increased cell eccentricity (0.74 vs 0.48, p <0.0001) and reduced variability in cell orientation (Δ = −47.1°, p <0.0001). At the transcriptome level, differential gene expression was extensive (continuous vs static: 2,103 genes; pulsatile vs static: 2,643 genes; pulsatile vs continuous: 384 genes). Continuous flow favoured oxidative-metabolic and barrier-maintenance pathways. By contrast, pulsatile shear of the same mean load promoted cell-cycle/checkpoint signalling rather than oxidative-metabolic programmes. Head-to-head, pulsatile flow emphasized MYC proto-oncogene (MYC) and E2F transcription factor (E2F), whereas continuous flow preferentially engaged oxidative phosphorylation and phosphoinositide 3-kinase (PI3K)–AKT serine/threonine kinase (AKT)–mechanistic target of rapamycin (MTOR) (with higher tumor protein p53and transforming growth factor-β signaling).

**Conclusion:** Under matched mean shear, pulsatile versus continuous flow drive distinct endothelial morphological and transcriptional programs, enriching pathways central to blood vessel function, remodelling, and disease. 3D macrofluidic platforms can replicate flow-specific endothelial cell mechanobiology, providing a translational tool to better understand vascular outcomes.

## INTRODUCTION

Endothelial cells (ECs) form a continuous, metabolically active monolayer with ∼10 trillion cells lining ∼350 m² of the adult vasculature [1, 2]. At the blood–tissue interface, ECs function as biochemical sensors and mechanotransducers, integrating circulating cues with hemodynamic forces to regulate anti-coagulant, anti-thrombotic, and vasoactive outputs, including nitric oxide, prostacyclin, endothelial-derived hyperpolarization, C-type natriuretic peptide, and membrane-bound constituents such as heparinoids, thrombomodulin, and tissue plasminogen activator [3, 4].

Within intact vessels, blood flow over complex geometries imposes spatially and temporally varying mechanical inputs on ECs including radial forces linked to intravascular volume, tangential forces at cell-cell interfaces, and the frictional forces of flowing blood - termed wall shear stress (WSS) [5]. In the arterial wall, WSS is the dominant endothelial stimulus [6], whereas vascular smooth muscle principally senses intraluminal pressure and circumferential strain [7]. As WSS correlates with viscosity, volumetric flow, and luminal radius, even minor changes due to curvature, taper, or branch points can generate large local changes in WSS that have a significant impact on endothelial phenotypes [8]. Endothelial function is therefore critically dependent on mechanotransduction by coupling luminal flow to nitric oxide (NO) production, ion-channel activity, cytoskeletal remodeling, gene expression, and ultimately tissue perfusion [5, 9]. While these core signalling pathways are conserved, venous ECs display distinct baseline set-points and sensitivity to shear that alter transcriptional and functional responses that are distinct from those in arterial ECs [10].

Endothelial mechanosensing relies on receptors and macromolecule complexes to initiate downstream signalling – these include integrins [11–13], receptor tyrosine kinases [14], G-protein–coupled receptors [15], ion channels [16, 17], and intercellular junctional proteins [18]. Flow magnitude and pattern also impact on distinct biologic states. Steady laminar WSS engages lipid signalling and kinase cascades that converge on phosphoinositide 3-kinase (PI3K)–AKT serine/threonine kinase (AKT)–dependent endothelial nitric oxide synthase (eNOS) activation, resulting in NO-mediated vasodilation and suppression of inflammatory programs [5]. Oscillating WSS increases expression of nuclear factor kappa-light-chain-enhancer of activated B cells (NF-κB), and promotes an adhesive, procoagulant, and pro-inflammatory phenotype typical of atherosclerosis-prone regions [9]. These shear-responsive signalling cascades control cytoskeletal architecture, polarity, junctional integrity, cell-cycle progression, and proliferation [19, 20]. Although the complete transcriptional control of mechanotransduction remains to be resolved, several key factors are involved, including Krüppel-like factor-2 (KLF2), NF-κB, activator protein-1 (AP-1), and Yes-associated protein (YAP) [21, 22].

While traditional two-dimensional (2D) cell culture provides compositional control and throughput, it constrains ECs to rigid 2D surfaces lacking physiological curvature and authentic hemodynamics. This results in a well-documented divergence in cell morphology, cell-cell junctional organization, proliferation, differentiation, metabolism, and pharmacologic responses relative to *in vivo* states [23–25]. Three-dimensional (3D) culture such as spheroids, organoids, and engineered tissues better recapitulate the morphology, polarity, junctional architecture, and transcriptome of ECs *in vivo* [26–30], but often do not recapitulate physiologic shear [26, 29]. Advances in high-resolution imaging and additive manufacturing now enable precise capture of patient-specific geometries [31, 32] and accurate fabrication [33, 34] that reproduces vessel curvature, taper, and branching. Growing endothelial cells on such printed constructs under controlled hemodynamic environments, generates a macrofluidic testbed that mimics the experimental control of classic *in vitro* systems, as well as the geometric determinants of WSS that govern endothelial phenotype *in vivo*.

The aim of this study was to develop a 3D-printed macrofluidic platform that supports endothelial cell growth, coupled with a pump system to enable perfusion of continuous laminar flow or programable, physiologically relevant pulsatile flow. The morphologic and transcriptomic impacts of these properties on ECs was characterised to demonstrate the critical role of both WSS and flow profile in endothelial mechanotransduction.

## METHODS

### Additive Manufacturing and PDMS Casting of Idealised Vessel Geometries

Idealized 6 cm vessels were designed using commercial CAD software, SolidWorks (2023; Dassault Systèmes, Vélizy-Villacoublay, France). Several soluble printing filaments were screened on a fused-deposition manufacturing 3D printer (Prusa i3 MK3S; Prusa Research, Prague, Czech Republic). These included 3 PVA filaments (eSUN PVA, 1.75 mm, Shenzhen eSUN Industrial Co., Ltd., Shenzhen, China; MatterHacker Pro Series, 1.75 mm, MatterHackers Inc., Lake Forest, CA, USA; FormFutura Aquasolve PVA, B.V., Nijmegen, The Netherlands), 1 BVOH filament (BASF Ultrafuse,, 1.75 mm, BASF 3D Printing Solutions GmbH, Heidelberg, Germany), and 1 lye-soluble copolymer (Xioneer VXL90,, 1.75 mm, Xioneer Systems GmbH, Vienna). These printing filaments were assessed using a temperature-tower calibration print (https://www.thingiverse.com/thing:2729076, Supplementary Figure 1). Post-print surface smoothing methods tested included: sandpaper, wet wiping, static dip in room temperature water for 30 s, static dip in room temperature water for 60 s, static dip in room temperature water for 180 s. Post-modification surface roughness was assessed using a scanning laser microscope (Keyence VK-X200, Keyence Corporation, Osaka, Japan) (Supplementary Figure 1). Polydimethylsiloxane (PDMS; Sylgard 184, Dow Corning, supplied by CBC Bearings, Australia) prepolymer and curing agent were mixed in a 10:1 weight ratio and degassed for 60 min. Once smoothed, PVA cores were positioned in acrylic resin molds (designed in commercial CAD software, SolidWorks) with a 0.5 mm offset and filled with PDMS. PDMS was cured at room temperature for 48 h and then immersed in agitated room-temperature water for 72 h to dissolve the PVA core geometries.

### ECM Surface Coating and Endothelial Monolayer Formation

Surface modification was tested by incubating PDMS surfaces in 1% (w/v) gelatin, 10 µg/mL fibronectin (Sigma-Aldrich, Merck; F0895-5MG), 20 µg/mL fibronectin, 50 µg/mL fibronectin, 1% (w/v) Pluronic F-127 (Invitrogen, Cat. P6866) for 1 h at 37°C (5% CO2) or by using a 1% (w/v) PEG-PDMS (Specific Polymers, Cat. SP-8P-3-001) additive to the PDMS prepolymer base agent (Supplementary Figure 2). Coating solution was removed and the surface air-dried for 60 min at room temperature. Human microvascular endothelial cells (HMEC-1, ATCC; CRL-3243) were maintained in MCDB-131 complete medium (Gibco; 10372019) supplemented with 10 ng/mL human epidermal growth factor (Sigma-Aldrich, Merck; E9644), 1 µg/mL hydrocortisone (Sigma-Aldrich, Merck; H-4001), 10 nM L-glutamine (Gibco; 25030-081), and 10% (v/v) fetal bovine serum (Gibco; 10099-141) at 37 °C and 5 % CO₂. HMEC-1 ECs were seeded onto the model surfaces at a density of 3.75 × 10⁴ cells/cm² in a sample spinner for 24 h at 37 °C and 5 % CO₂ (Supplementary Figure 3).

### Manufacturing of Flow System and Generation of Flow

3D components were designed using Solidworks CAD software and fabricated using heat-resistant resin (TR-300 resin, Phrozen, Hsinchu, Taiwan; Cat. No. TR-300) on a stereolithography 3D printer (Sonic Mighty 12K SLA printer, Phrozen, Hsinchu, Taiwan). The perfusion system consisted of a closed loop incorporating two 100 mL MACS GMP Cell Differentiation Bags (Miltenyi Biotec, Bergisch Gladbach, Germany; Cat. No. 170-076-400) as culture media reservoirs, custom-printed connectors, and peristaltic pump silicone tubing LongerPump #25 tubing, LongerPump Co., Ltd., China). Flow was generated using a programmable peristaltic pump (NE-9000; New Era Pump Systems Inc., Farmingdale, NY, USA) and monitored using an ultrasonic flow meter (Sonoflow CO.55; Sonotec GmbH, Halle, Germany) positioned immediately upstream of the vessel segment. To generate pulsatile flow, patient-derived Doppler ultrasound data from arteriovenous fistulas (n = 13) were analyzed to extract magnitude of baseline flow, time at baseline flow, time to peak flow, magnitude of peak flow, time at peak flow, time to post-peak baseline, magnitude of post-peak baseline flow, and time at post-peak baseline flow over one cardiac cycle. Across the cohort, the mean volumetric flow was 595 ± 281 mL/min, with an average minimum of 463 ± 240 mL/min and an average maximum of 779 ± 352 mL/min. The average cardiac cycle duration was 0.84 s, with mean time to peak 0.334 s, peak duration 0.20 s, and time from peak back to baseline 0.347 s. The flow profiles from these data were scaled to achieve an average flow of 100 mL/min (85-145 mL/min over 0.84 s) and programmed to generate a realistic pulsatile waveform. Flow conditions were maintained for 24 h of perfusion following an initial 24 h static attachment phase. Except for the pump, the system was housed and operated within a humidified incubator at 37°C (5 % CO₂).

Flow characteristics under continuous flow and pulsatile flow conditions were approximated under the assumptions of steady laminar flow in a straight, smooth, rigid, circular tube of 4.8 mm internal diameter. The culture medium was modelled as an incompressible, Newtonian, isothermal fluid with constant viscosity and density. Gravitational and entrance effects, wall compliance, and fluid – structure interaction were neglected. With these assumptions, hemodynamics follow the bulk-flow law [35] where flow pattern is characterized by the Reynolds number (Re) = ρVD/μ (ρ: density; μ: viscosity; V: bulk velocity; D: diameter). WSS (τw) for laminar tube flow is given by τw = 4μQ/(πr³), or equivalently τw = 8μV/D. Pulsatility index (PI) was defined as PI = (Qmax – Qmin)/Qmean, and for pulsatile flow, unsteadiness was quantified using the Womersley number (α), defined as α = R(ω/ν)½ (R: radius; ω: angular frequency; ν: kinematic viscosity). For pulsatile conditions, a quasi-steady approximation was applied such that instantaneous wall shear stress scaled linearly with the instantaneous flow rate with no flow reversal.

### Morphological assessment by fluorescence microscopy

At the end of flow exposure, the constructs were fixed and stained with Phalloidin–Atto-565 (94072; Merck, Darmstadt, Germany) and counterstained with DAPI (D1306; Thermo Fisher Scientific, Waltham, MA, USA). Imaging was performed using a Zeiss LSM 880 confocal microscope (Carl Zeiss, Oberkochen, Germany) housed at the Katharina Gaus Light Microscopy Facility, Mark Wainright Analytical Centre, UNSW Sydney. Images were analyzed in Fiji (ImageJ) (Fiji distribution v2.16.0 / ImageJ v1.54p; RRID:SCR_002285 / RRID:SCR_003070) with default settings.

### Bulk RNA Sequencing

Total RNA was isolated using a RNeasy Plus Mini Kit (74134, QIAGEN, Hilden, Germany) according to manufacturer’s instructions. RNA integrity was verified using an Agilent TapeStation system (Agilent Technologies, Santa Clara, CA, USA), with all samples showing excellent quality (RINe ≥ 9.9) (Supplementary Figure 4). Libraries were constructed from total RNA using the Illumina stranded mRNA workflow, dual-indexed, PCR-amplified within recommended cycle limits, pooled equimolarly, and sequenced on an Illumina NovaSeq X by the Ramaciotti Centre for Genomics (UNSW Sydney, Australia) (Supplemental Figure 5). Raw paired-end RNA-seq reads were staged from archives and assessed with FastQC for base quality, Q30 rate, GC content, adapter/contaminant sequences, and duplication. Reads were adapter/quality trimmed with fastp where possible, and FastQC/fastp outputs were aggregated with MultiQC. Trimmed reads were aligned to the human genome using Rsubread, BAMs were QC-checked, and gene-level counts were generated with featureCounts. Differential expression was performed in DESeq2 with apeglm shrinkage [36] of log₂ fold changes and significance defined as adjusted p < 0.05 and |log₂FC| ≥ 1. Hallmark pathway enrichment was assessed by over-representation (clusterProfiler) and preranked GSEA (fgsea). Results were ranked by false discovery rate (FDR) and normalized enrichment score (NES). The top 30 enriched pathways (FDR < 0.05) are reported. The full pipeline is summarized in Figure 1.

**Figure 1:**
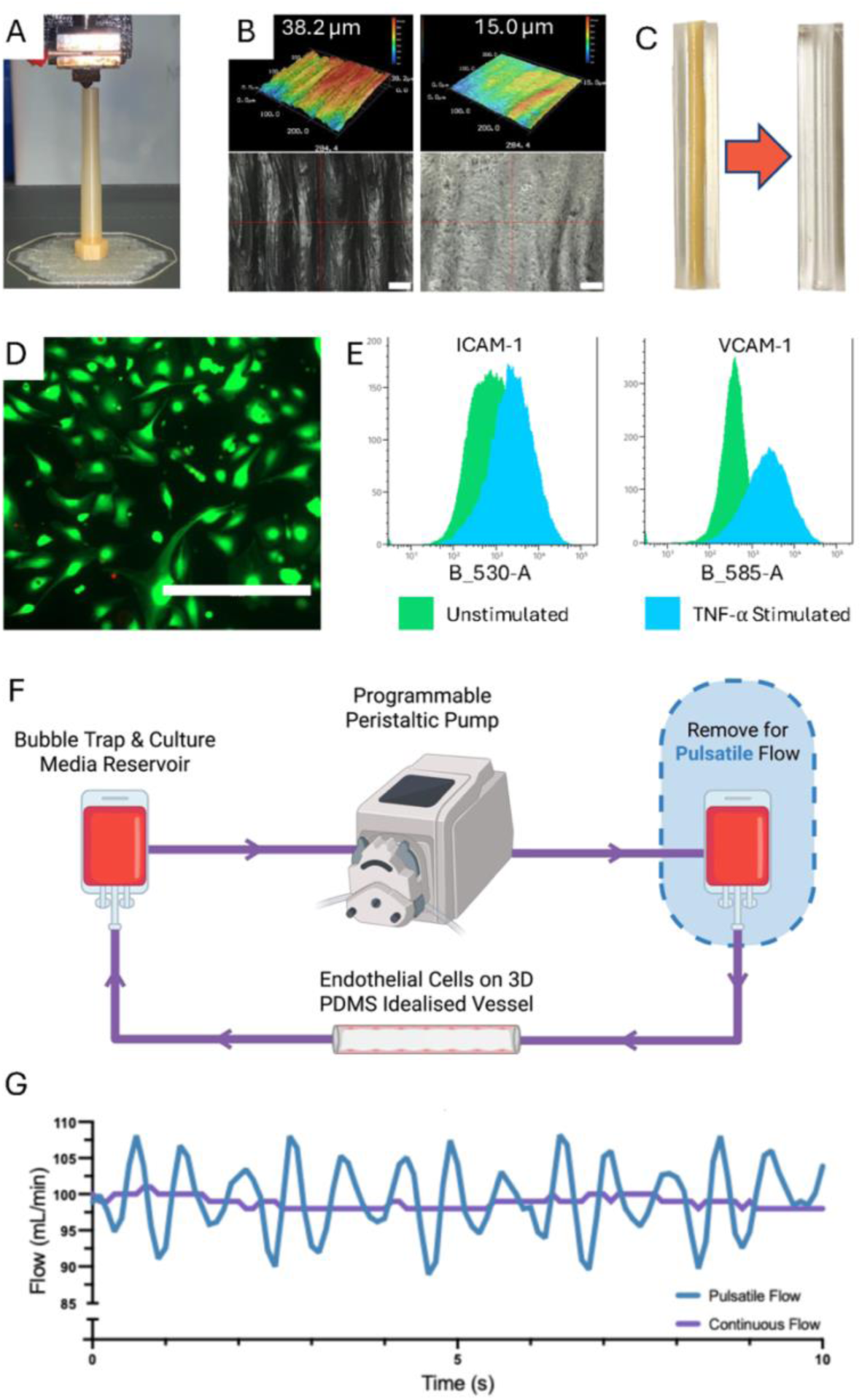
Fabrication and validation of a 3D vessel-mimetic flow platform. (**A**) 3D-printed idealized vessel used as the sacrificial core/master for casting. (**B**) Surface smoothing optimisation was assessed by laser profilometry. A 30 s water dip was found to be the most effective method. The numeric labels above each panel indicate the measured mean roughness (µm) for the imaged field. (**C**) PDMS casting around the printed core and subsequent dissolution of the core to yield a perfusable channel. (**D**) HMEC-1 viability on fibronectin coated PDMS was confirmed by Live/Dead staining. (**E**) Flow cytometry validation of endothelial responsiveness on PDMS: 6 h TNF-α stimulation increases ICAM-1 (CD54) and VCAM-1 (CD106) expression compared with unstimulated controls. (**F**) Schematic of the closed-loop perfusion rig showing bubble trap/media reservoir and a programmable peristaltic pump. The second bubble trap (immediately downstream of peristaltic pump) can be removed when modelling physiologic flow. (**G**) This setup delivers either steady continuous flow (purple) or a pulsatile (blue) waveform as demonstrated by a representative trace.

### Statistical Analysis

Descriptive statistics are reported as frequencies (%) for categorical variables, mean ± SD for continuous normally distributed variables, and median (IQR) for non-normal continuous variables. Group differences were tested with one-way ANOVA. If variances were unequal, Welch’s ANOVA was used. For non-normal data, the Kruskal-Wallis test substituted for ANOVA. Post-hoc testing after ANOVA employed Tukey’s honestly significant difference for all pairwise comparisons. For image-derived morphology cell-level measurements were aggregated to a sample-level. A p value < 0.05 was considered statistically significant. All analyses for non-sequencing data were performed in GraphPad Prism Version 10.3.1 (464; 21 Aug 2024).

## RESULTS

### Construction and Characterisation of a 3D Macrofluidic System

Soluble printing filaments were screened using temperature-tower calibration prints. The most consistent extrusion and dimensional fidelity was achieved with eSUN PVA (Supplementary Figure 1A). Trials of surface smoothing methods (Supplementary Figure 1 B-G) identified a 30 s static water dip reduced topographic variability (47.4 +/- 14.8 µm to 14.03 +/- 0.9 µm, p <0.001) without introducing deforming features (Figure 2B). PVA prints were cast in PDMS then washed away once the PDMS was cured – leaving behind a lumen within the PDMS of the printed geometry (Figure 2C). Among multiple surface coatings tested (Supplementary Figure 2), a single-component fibronectin coating at a concentration of 10μg/mL for between 1 and 24 h provided the most robust cellular adhesion. Seeding HMEC-1 cells at a starting density of 3.75×10^4^ cells/cm^2^ consistently produced a confluent layer of endothelial on the PDMS lumen after 24 h (Supplementary Figure 3). The HMEC-1 monolayer protocol derived from these data included fibronectin surface modification between 1–24 h, HMEC-1 seeding density 3.75×10⁴ cells/cm², and culture in static conditions for ≥24 h to obtain a robust endothelial monolayer. Live/Dead staining confirmed viability of all adherent cells under these conditions (Figure 2D). Preservation of EC functional responsiveness was confirmed, with TNF-α stimulation resulting in significant increases in ICAM-1 (272.3 ± 23.0%, p < 0.001) and VCAM-1 (648.2 ± 66.6%, p < 0.01) expression compared to controls (Figure 2E). Continuous flow was generated with a mean flow of 98.85 ± 0.8 mL/min (minimum 98 mL/min, maximum 101 mL/min) (Figure 2G). Pulsatile flow was modelled from patient arteriovenous fistula doppler ultrasounds to generate a mean flow of 99.32 ± 5.66mL/min (minimum 85, maximum 110) over a 0.84s pulse cycle (Figure 2F).

**Figure 2 -.**
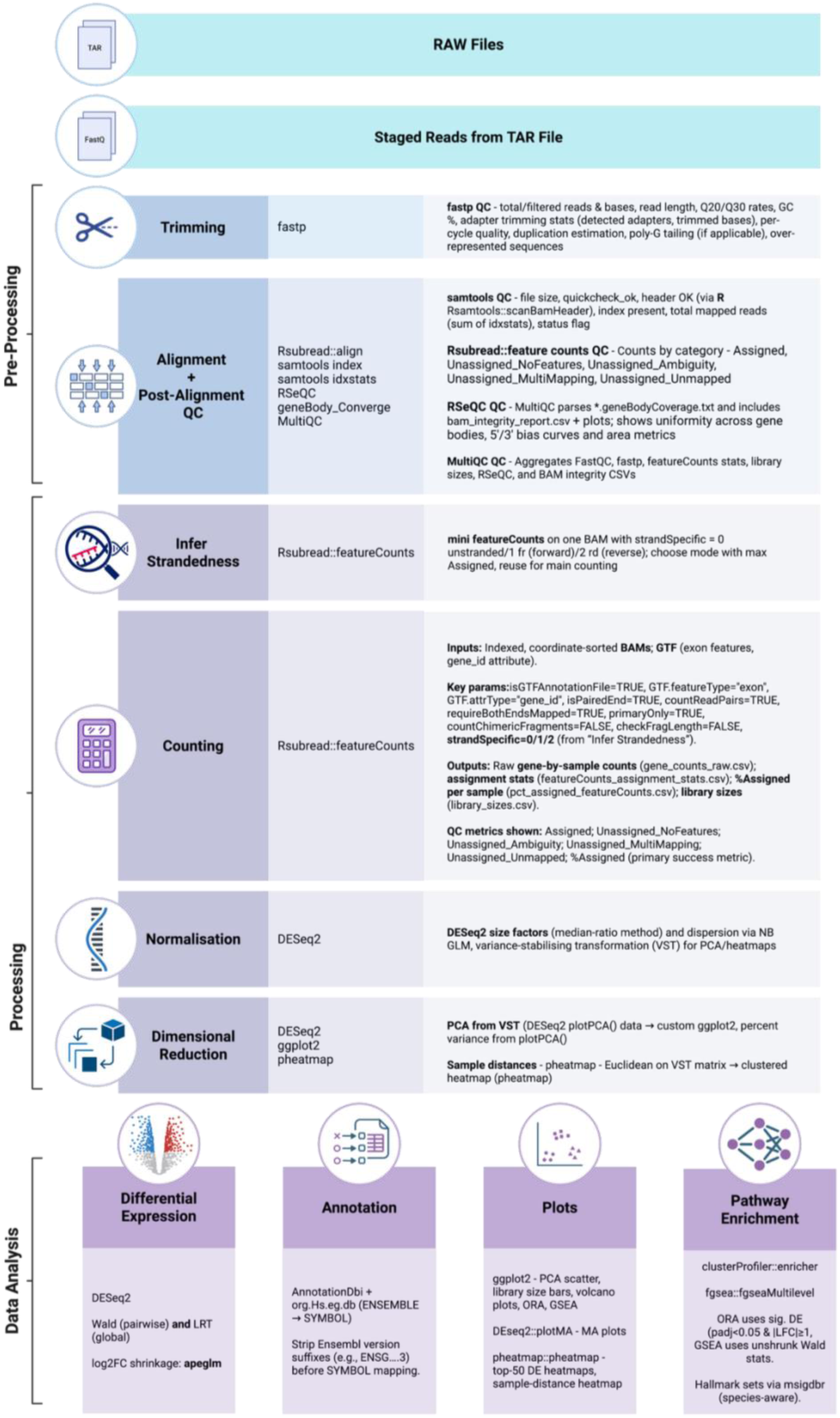
Bulk RNA-seq workflow overview. QC = quality control, Q20/Q30 = Phred quality thresholds, GC% = guanine–cytosine content, BAM = binary alignment/map, PCA = principal component analysis, VT = variance-stabilizing transform, NB GLM = negative-binomial generalized linear model, DE = differential expression, log2FC = log2 fold change, LRT = likelihood-ratio test, ORA = over-representation analysis, GSEA = gene set enrichment analysis, HGNC = HUGO Gene Nomenclature Committee.

With diameter D = 4.8 mm, the continuous flow profile yielded *τ*w of 0.150 Pa (1.50 dyn/cm²) at trough, 0.152 Pa (1.52 dyn/cm²) at mean, and 0.155 Pa (1.55 dyn/cm²) at peak, with corresponding *Re* of 433, 437, and 447, respectively. The pulsatile flow profile spanned *τ*w 0.130 – 0.169 Pa (1.30–1.69 dyn/cm²) from trough to peak (mean 0.153 Pa, 1.52 dyn/cm²) and Re 376 – 486 (mean 439). In both cases, wall shear stress remained positive along the conduit, with no inlet or localised flow reversal throughout the cycle owing to the idealised vessel geometry, yielding an oscillatory shear index (OSI) of 0. For the continuous flow condition, Q_mean_ = 98.85 mL/min, Q_min_ = 98 mL/min and Q_max_ = 101 mL/min, making PI_cont_ ≈ 0.03. For the pulsatile condition, Qmean = 99.32 mL/min, Qmin = 85 mL/min and Qmax = 110 mL/min, giving PIpulse ≈ 0.25. As wall shear stress scales linearly with volumetric flow under these laminar conditions, the shear-based PI was effectively identical to the corresponding flow-based PI in each condition. Womersley number (α) was calculated as α = Rω/ν, where R = D/2 = 2.4 × 10⁻³ m, the cardiac cycle T = 0.84 s gave a frequency f = 1/T ≈ 1.19 Hz, ω = 2πf ≈ 7.5 rad/s, and the kinematic viscosity of the medium was ν ≈ 1.0 × 10⁻⁶ m²/s, yielding α ≈ 6.6.

### Flow Impacts Endothelial Cell Morphology

Endothelial morphology and cytoskeletal organization varied as a function of flow as measured by immunofluorescent staining (Figure 3). Under static conditions, cells displayed a cobblestone morphology with no preferred orientation (Figure 3A-B). After 24 h of continuous or pulsatile flow, cells elongated and F-actin bundles aligned with the direction of flow (Figure 3 C-D). Quantification confirmed greater eccentricity under both flow regimens compared to static conditions (0.74 ± 0.16 in continuous flow, 0.60 ± 0.18 in pulsatile flow, 0.48 ± 0.11 in static conditions, p <0.0001 continuous flow vs. static, p =0.004 pulsatile flow vs. static culture) (Figure 3G). Orientation variability was highest under static conditions (56.63° ± 41.42°), whereas exposure to flow reduced variability, with significantly lower dispersion under continuous compared with pulsatile flow (9.484° ± 8.11° vs 33.70° ± 15.71°, p = 0.03) (Figure 3H).

**Figure 3:**
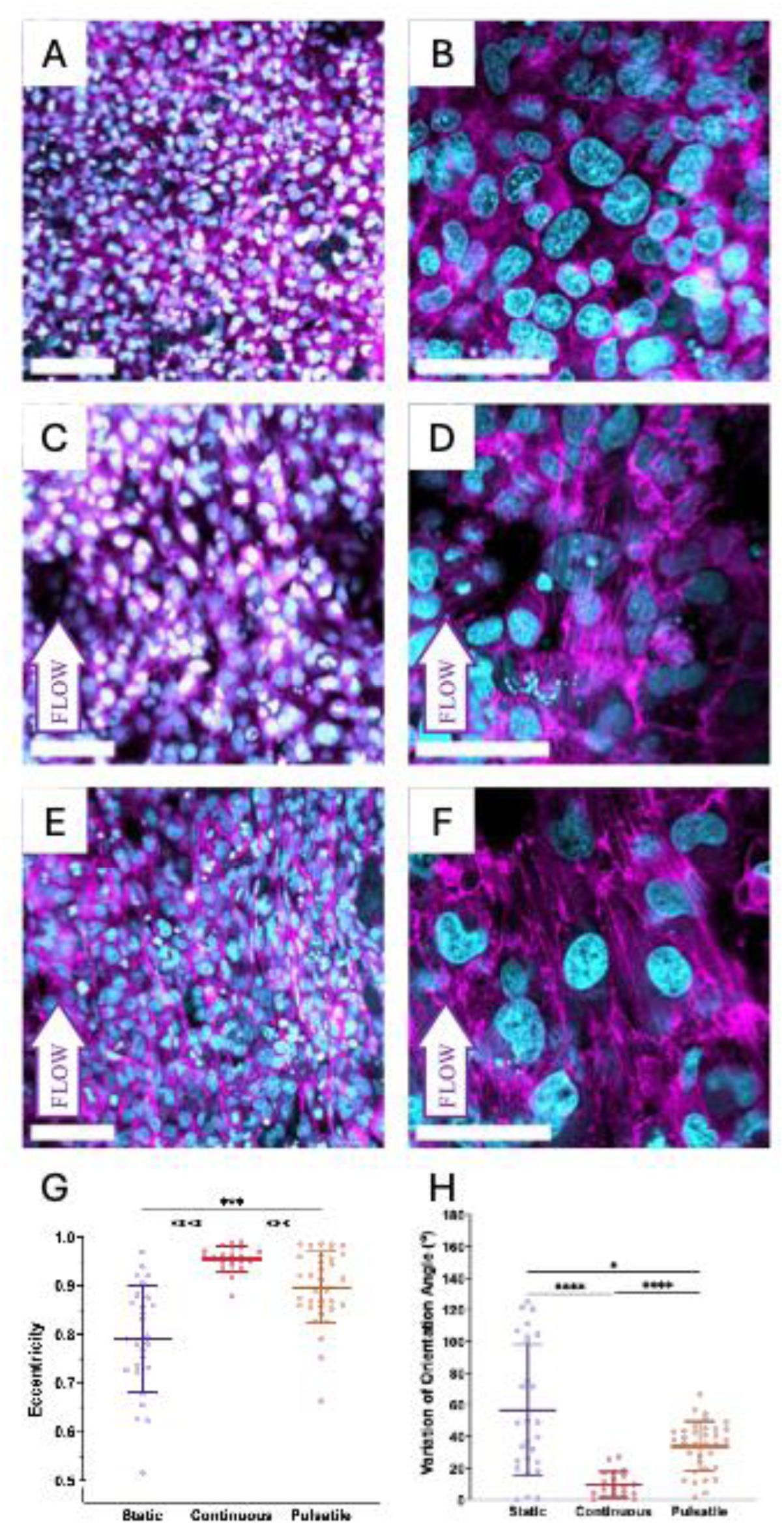
Flow-dependent remodeling of cell morphology and cytoskeleton. HMEC-1 cells were grown on fibronectin coated PDMS macrofluidic models and then exposed to flow or control conditions for 24 h. (**A, B**) Static culture for 24 h. (**C, D**) Continuous flow for 24 h. (**E, F**) Pulsatile flow for 24 h. Cyan, nuclei (DAPI); magenta, F-actin (phalloidin-Atto565). (**G**) Quantification of cell eccentricity. (**H**) Variation in orientation angle. Scale bars: 100 μm (**A, C, E**) or 50 μm (**B, D, F**). Each point marks a cell. * = p < 0.05, *** = p<0.001, **** = p<0.00001.

### Flow Alters Transcription of Shear-Responsive Genes

Bulk RNA sequencing was performed on three independent biological replicates per condition. Principal component analysis demonstrated clustering of replicates (Figure 4A).

**Figure 4:**
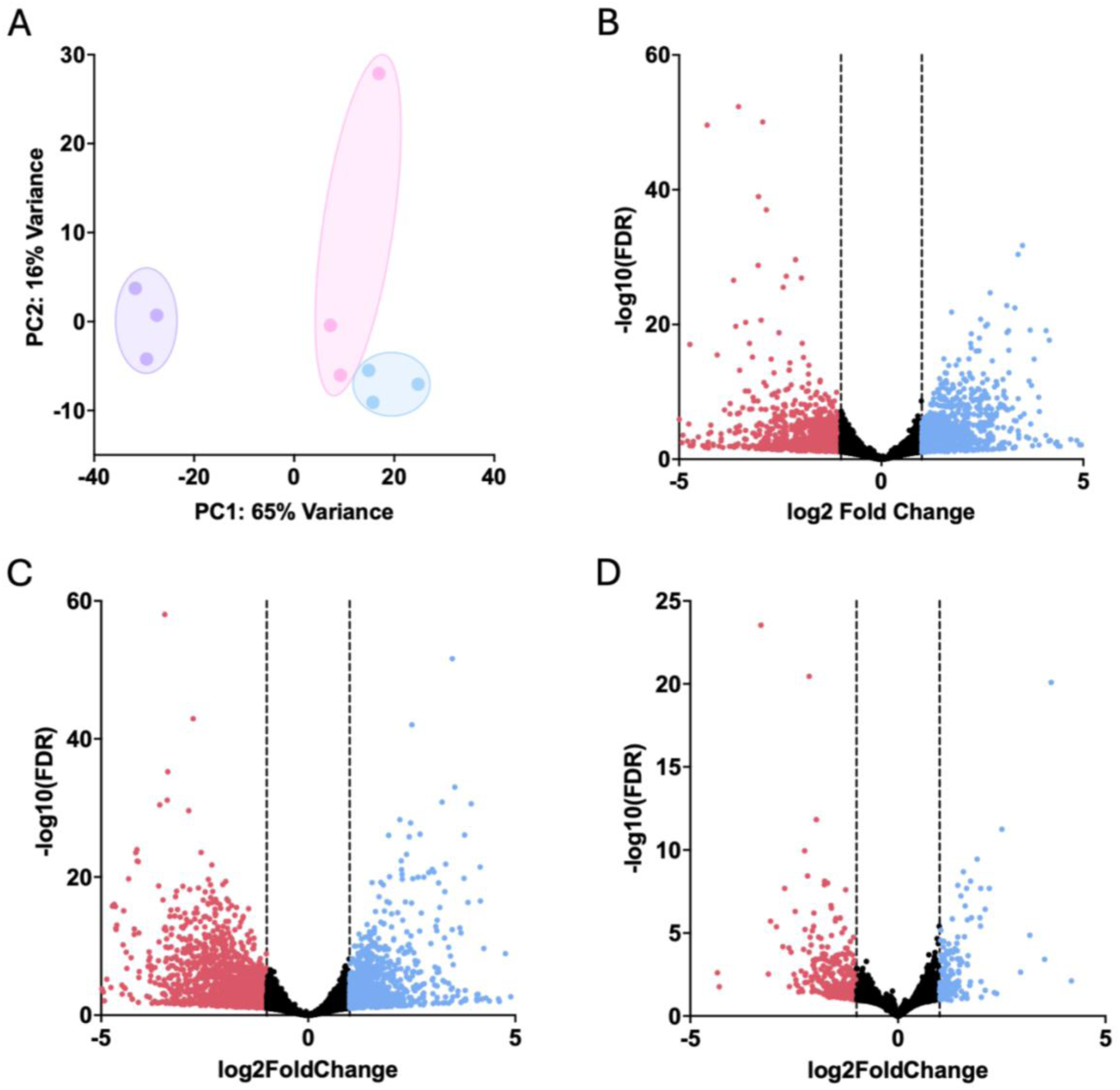
Altering flow profile results in changes in transcription. (**A**) Principal component analysis (PCA) of variance-stabilized counts (n=3 biological replicates per condition) demonstrated separation of cells maintained under static conditions (purple) from flow-exposed samples and a secondary separation between continuous (pink) and pulsatile (blue) flow. (**B–D**) Volcano plots depict differential expression (DESeq2; coloured points denote padj < 0.05 with |log_2_FC| ≥ 1; dashed lines mark ±1 log_2_FC). (**B**) Continuous flow vs. static, (**C**) Pulsatile flow vs. static. (**D**) Pulsatile vs. continuous flow.

In the continuous flow vs. static condition comparison, a total of 2,103 genes met the significance criteria, with 856 increased and 1,247 decreased under continuous flow. Relative to static culture, pulsatile flow similarly induced a canonical shear response. Under pulsatile flow a total of 2,643 genes were significantly differentially expressed, with 1,017 increased and 1,626 decreased (Figure 4B). A heat map of the top 50 differentially expressed genes comparing continuous flow with static culture recapitulated the PCA analysis, with obvious clustering of flow conditions compared to static culture (Figure 5A).

**Figure 5:**
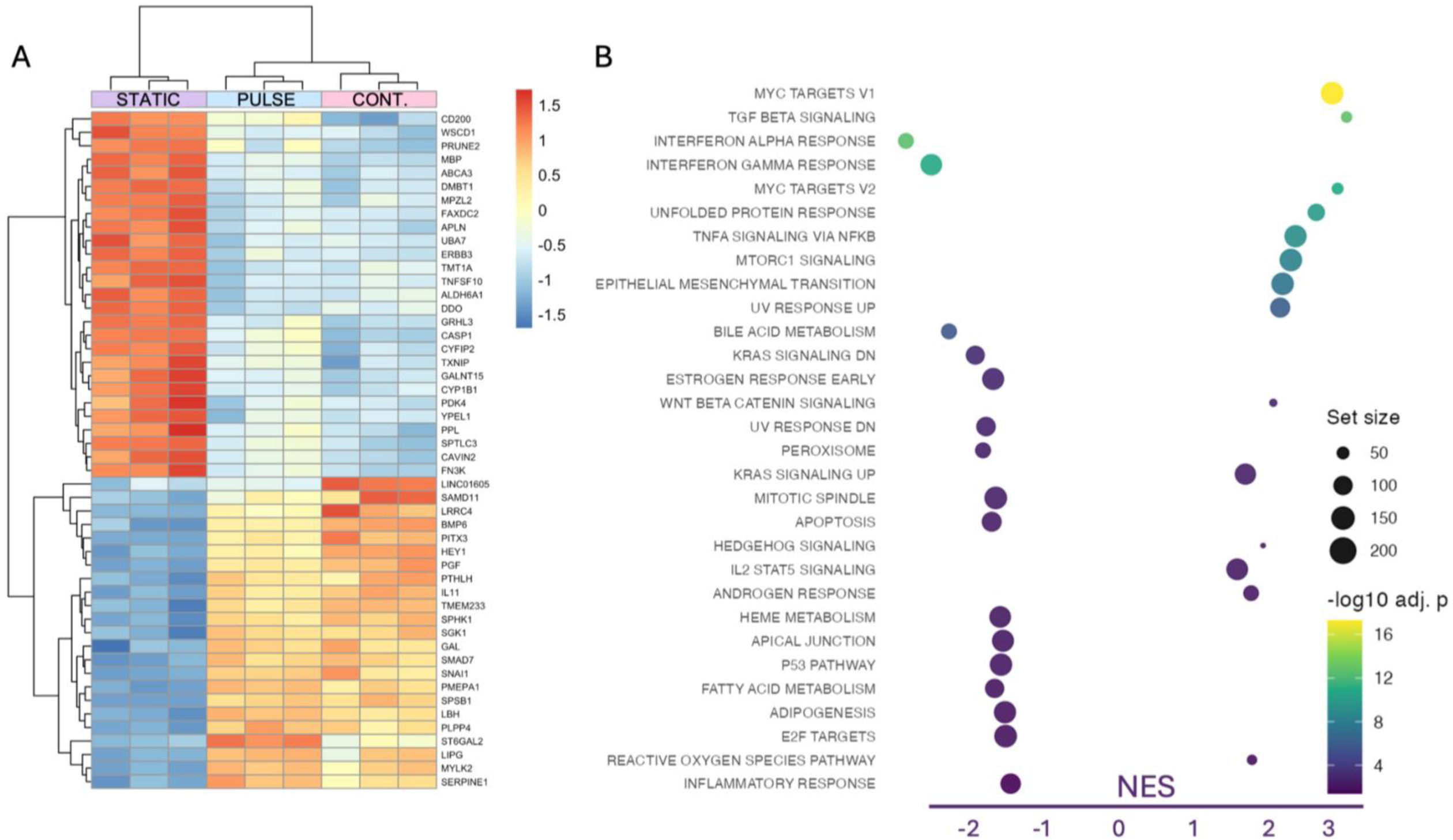
Heat map of the top 50 differentially expressed genes in continuous flow vs. static conditions comparison across all samples (columns). Values are variance-stabilized counts (DESeq2) and z-scored by gene. Rows and columns were hierarchically clustered using 1–Pearson correlation distance and complete linkage. Panel B shows normalised enrichment score (NES) for Pulsatile vs Continuous flow (positive = enriched in pulsatile; negative = enriched in continuous).

### Flow Suppresses Inflammation and Proliferation but Enriches Junctional-Metabolic Signalling

Pre-ranked gene set enrichment analysis of Hallmark pathways for continuous flow versus static showed positive enrichment in continuous flow for MYC targets V1 (NES = 2.66, P = 5.11×10⁻¹⁸), TGF-β signalling (NES = 2.84, P = 1.78×10⁻¹³), MYC targets V2 (NES = 2.73, P = 1.10×10⁻¹¹), unfolded protein response (NES = 2.46, P = 1.02×10⁻¹⁰), TNFα signalling via NF-κB (NES = 2.21, P = 6.58×10⁻¹⁰), mTORC1 signalling (NES = 2.15,P = 4.89×10⁻⁹), and epithelial-mesenchymal transition (NES = 2.04, P = 3.50×10⁻⁸). Negative enrichment included interferon-γ response (NES = −2.34, P = 1.10×10⁻¹¹), interferon-α response (NES = −2.20, P = 1.82×10⁻¹⁰), UV response up (NES = −1.93, P = 1.37×10⁻⁶), bile-acid metabolism (NES = −2.12, P = 1.78×10⁻⁶), KRAS signalling DN (NES = −1.79, P = 7.90×10⁻⁴), and peroxisome (NES = −1.69, P = 2.84×10⁻³) (Figure 5).

For pulsatile flow versus static conditions, positively enriched pathways in pulsatile flow included MYC targets V1 (NES = 2.61, P = 2.38×10⁻²⁵), interferon-α response (NES = 2.16, P = 1.26×10⁻¹⁴), interferon-γ response (NES = 1.98, P = 3.29×10⁻¹⁰), epithelial–mesenchymal transition (NES = 1.96, P = 3.92×10⁻⁷), unfolded protein response (NES = 1.77, P = 1.82×10⁻⁶), and mTORC1 signalling (NES = 1.66, P = 1.94×10⁻⁴). Negative enrichment was observed for adipogenesis (NES = −2.01, P = 6.47×10⁻¹²), inflammatory response (NES = −1.91, P = 1.33×10⁻⁹), apical junction (NES = −1.90, P = 1.33×10⁻⁹), estrogen response early (NES = −1.90, P = 1.92×10⁻⁹), WNT/β-catenin signalling (NES = −1.79, P = 4.29×10⁻⁸), and fatty-acid metabolism (NES = −1.36, P = 4.35×10⁻³) (Supplemental Figure 6).

### Flow Pattern Alters Transcription of Shear-Responsive Genes

In a direct comparison of pulsatile versus continuous flow, 149 genes were upregulated and 235 genes were downregulated during pulsatile flow compared to continuous flow.

Positive enrichment in pulsatile flow included oxidative phosphorylation (NES = 2.19, P = 1.05×10⁻⁵), UV response up (NES = 1.83, P = 1.64×10⁻⁴), epithelial–mesenchymal transition (NES = 1.59, P = 2.00×10⁻³), and coagulation (NES = 1.58, P = 2.00×10⁻³). Negative enrichment included mitotic spindle (NES = −2.17, P = 1.07×10⁻²³), UV response down (NES = −2.12, P = 4.59×10⁻¹⁸), G2M checkpoint (NES = −2.03, P = 5.64×10⁻¹⁵), MYC targets V2 (NES = −2.00, P = 1.84×10⁻¹²), E2F targets (NES = −1.88, P = 2.29×10⁻¹⁰), P53 pathway (NES = −1.83, P = 1.67×10⁻¹⁰), and TGF-β signalling (NES = −2.05, P = 1.13×10⁻⁹) (Supplemental Figure 7).

A heat map of the top 50 differentially expressed genes comparing pulsatile flow with continuous flow reinforced these differences and recapitulated the PCA structure clearly showing the separation under the differing flow conditions (Supplemental Figure 7).

## DISCUSSION

We engineered a vessel-mimetic, 3D-printed macrofluidic system that reproduces continuous and physiologic pulsatile waveforms, providing a controllable testbed to separate and identify how the presence and temporal structure of flow programs endothelial cell phenotype and function. Static culture, continuous flow, and pulsatile flow generated distinct endothelial transcriptomes. Culture under static conditions was associated with increased cell circularity, elevated adhesion molecule expression, and enrichment of interferon and inflammatory signalling relative to cells exposed to flow. Continuous unidirectional shear showed relative enrichment of oxidative phosphorylation and PI3K–AKT–mTOR signalling, consistent with redox homeostasis and adaptive stress control, whereas pulsatile shear did not. Pulsatile shear with the same mean load yielded an intermediate morphology and higher expression of adhesion-related transcripts than continuous flow, and more strongly enriched cell-cycle and checkpoint pathways. Collectively, these observations validate the macrofluidic model as a physiologically informative platform and highlight that endothelial cells are exquisitely sensitive to the temporal structure of flow.

In this system, continuous and pulsatile profiles were closely matched for mean flow, Reynolds number, and time-averaged WSS (OSI ≈ 0; no flow reversal). Nevertheless, waveform shape remained a key determinant under constant mean WSS. This divergence despite matched means indicates sensitivity to features such as oscillatory shear index and the rate of change of shear. The pulsatile flow conditions showed a markedly higher PI (PI_pulse_ ≈ 0.25), and a Womersley number α ≈ 6.6, consistent with physiologically relevant, large-artery–like pulsatility. These antagonistic behaviours parallel our finding that smooth, unidirectional laminar shear best enforces endothelial cell quiescence, whereas pulsatility biases the cells toward proliferation and checkpoint programs, even without reversal of flow. At the pathway level, continuous and pulsatile flow profiles diverged from what weas observed for cells maintained in static culture but with distinct emphasis. Continuous flow displayed relative enrichment of interferon-α and TNFα–NF-κB signaling as well as oxidative phosphorylation and PI3K–AKT–mTOR signaling but did not activate MYC/E2F signalling. This is a profile that, despite higher inflammatory signaling than what was observed under pulsatile flow conditions, is more quiescent overall as evidenced by suppression of MYC/E2F. In contrast, pulsatile flow positively enriched MYC/E2F cell-cycle programs (MYC TARGETS V1/V2, E2F TARGETS, G2M CHECKPOINT) and UV-response signatures, and reduced oxidative phosphorylation, PI3K–AKT–mTOR, TNFα, NF-κB and interferon-α pathways at matched means. This signalling pattern indicates that pulsatile flow preferentially engages cell-cycle and checkpoint programs without the metabolic upshifts seen under continuous shear, with accompanying cytoskeletal plasticity that can facilitate remodeling. In contrast, suppression of cell-cycle and checkpoint pathways under continuous flow suggests tighter control of proliferation and reinforcing endothelial homeostasis. Under continuous shear, HMEC-1 cells adopted a quiescent, barrier-stabilizing profile, whereas modest pulsatility did not augment oxidative phosphorylation or PI3K–AKT–mTOR; instead pulsatility preferentially engaged MYC/E2F/G2M and UV-response programs, collectively defining a state primed for cell-cycle progression and checkpoint control.

These differences align with established mechanobiology in which waveform features govern endothelial state, rather than mean load alone. In landmark experiments that reconstructed the human carotid artery bifurcation hemodynamics and replayed region-specific waveforms in vitro, irregular oscillatory profiles activated NF-κB, increased expression IL-8, and heightened cytokine-inducible adhesion molecules, while steady laminar profiles upregulated KLF2, increased NO signalling, and stabilized junctions [37–39]. In the human carotid sinus, high oscillatory shear index at low mean shear arises from cardiac-cycle separation-boundary motion, yielding pro-inflammatory mechanical inputs [40]. Pathologies dominated by disturbed shear including the epicardial coronaries, carotid bifurcation/sinus, and peripheral conduit arteries are classic settings in which waveform shape governs endothelial state and disease expression. Our data therefore speak to mechanisms relevant to atheroprone vascular beds where oscillatory flow prevails. Through optimization of blood pressure and heart rate, modifying surgical or endovascular geometry, or tailoring device design and haemodynamic targets, it may be possible to shift endothelial phenotype toward a more quiescent, atheroprotective state even when mean flow and average shear are unchanged.

Prior work reproducing patient-specific waveforms derived from MRI and ultrasound-informed CFD with computer-controlled devices has focused on emphasizing accurate magnitude, frequency, and shape [41]. Our data have extended this by demonstrating that even when mean wall shear stress, Reynolds number and time-averaged metrics are held constant, differences in waveform shape alone drives opposing signalling pathways across proliferation, metabolism, junctional composition, and inflammatory tone. this implies that endothelial cells are sensitive to subtleties in flow such as the rate of change of shear stress over time. This observation captures the complex interplay between force magnitude, directional stability, rise and decay kinetics, and duty cycle . These waveform features are sensed and integrated by the endothelial glycocalyx, integrins, ion channels, junctional complexes, and YAP–TAZ mechanotransducers, which together regulation of NO bioavailability, redox balance, and transcriptional control [5, 18, 20, 21]. The findings described in this study demonstrate that even in a controlled in vitro system, small changes to a single factor can alter gene expression and phenotype in ways that can alter endothelial cell function. These results highlight the importance of explicit reporting of waveform descriptors including mean wall shear stress, oscillatory shear index, pulsatility amplitude and frequency, and rise and decay times, to ensure reproducibility, experimental fidelity, and external validity in studies of flow–function relationships.

Endothelial cell vascular bed of origin may also modulate the thresholds at which flow alters phenotype. Although core shear-responsive factors such as KLF2, KLF4, and NOS3 are induced in both human umbilical vein endothelial cells (HUVEC) and human aortic endothelial cells (HAEC), pathway outputs spanning cell cycle, oxidative stress, TGF-β signaling, angiogenesis, and sentinel genes including VCAM1, ICAM1, SELE, NOX4, and SOD2 display bed-specific changes or opposite trajectories, indicating lineage-encoded thresholds [10]. Maurya et al., showed that HAECs exposed to shear stress mount a faster and broader early response with more differentially expressed genes at 1-4 h and show distinct regulation of KLF4, E2F1, and inflammatory mediators relative to HUVECs [10]. In another study, exposure of HUVECs, HPAECs, and HMVECs to matched laminar shear increased canonical mechanotransduction in all cell types[42]. Despite this, only 44.2% of the differentially expressed genes were shared and HMVECs remained transcriptionally distinct by principal component analysis, indicating lineage-gated decoding of shear signals [42]. Accordingly, endothelial subtype and waveform should be treated as co-determinants of state rather than interchangeable experimental details.

In the present study we used the HMEC-1 EC cell line, acknowledging that their metabolic and mechanosensory set points are similar to venous ECs. Future studies should extend these analyses to primary human arterial and venous endothelial cells, including patient-derived cells, to test generalizability of the current findings across vascular beds and to compare responses in other established cell lines versus primary cultures. With further refinement, vessel-mimetic 3D macrofluidic platforms that deliver validated flow waveforms will provide an opportunity to improve mechanistic fidelity, reproducibility, and translational relevance. By matching device geometry and material compliance to patient -specific morphologies and by enforcing verified waveforms, such systems can preserve the hemodynamic conditions required for endothelial polarization, glycocalyx integrity, junctional architecture, and NO signalling that cannot be achieved in conventional 2D culture. Translational relevance would further increase by coupling these platforms to computational fluid dynamics derived from clinical imaging, permitting imposition of patient-level waveforms such as those encountered in arteriovenous fistulas or pre-/post-stenotic segments, and by aligning readouts with clinical endpoints (vasodilator reserve, permeability, leukocyte adhesion, thrombogenicity, anti-restenotic responses). The approach is inherently adaptable to vessel class and patient morphology. Channel geometry, compliance, branch angle, curvature, and extracellular matrix can be engineered to model arterial versus venous beds, high-flow states, or disturbed-flow niches. Integration of the platform that is described in the present study with patient-derived cells (primary ECs/SMCs or induced pluripotent stem cell derivatives) would enable a personalized medicine pipeline to be developed in which devices configured from an individual’s CT angiography or duplex ultrasound could be used to test medical devices, drug-elution strategies, or flow-reduction procedures under patient-specific hemodynamic conditions. A practical workflow for this would comprise imaging and segmentation, computational fluid dynamics, device fabrication, waveform calibration with inline pressure and flow sensors, patient-derived endothelial seeding, and exposure to the corresponding wall shear profiles, with orthogonal transcriptomic and functional readouts to verify mechanistic targeting. While a fully integrated pipeline of patient-specific geometry, patient-derived cells, and closed-loop waveform control remains to be realized, the present study establishes a reproducible, mechanistically faithful framework for endothelial modelling.

## AUTHOR CONTRIBUTIONS

NAS – conceived the topic, designed the experiments, performed laboratory work, collated all results, performed all analyses, and drafted the manuscript.

KAR – critically appraised the manuscript.

ZHE – developed the topic, supervised the research, and critically appraised the manuscript. TJB – developed the topic, supervised the research, and critically appraised the manuscript. JHE – developed the topic, supervised the research, and critically appraised the manuscript. BJC – conceived the topic, supervised the research, and critically appraised the manuscript.

## CONSENT FOR PUBLICATION

Not applicable.

## AVAILABILITY OF DATA AND MATERIALS

Raw data and full analysis code are not posted publicly. Processed results (normalized counts, differential expression summaries, and gene-set enrichment statistics) are included in the Supplementary Materials. Additional materials (selected intermediate files and executable notebooks) will be shared by the corresponding author upon reasonable request under a data use agreement. All software, versions, and parameters are specified in Methods.

## SUPPLEMENTAL FIGURES & TABLES

**Supplementary Figure 1:**
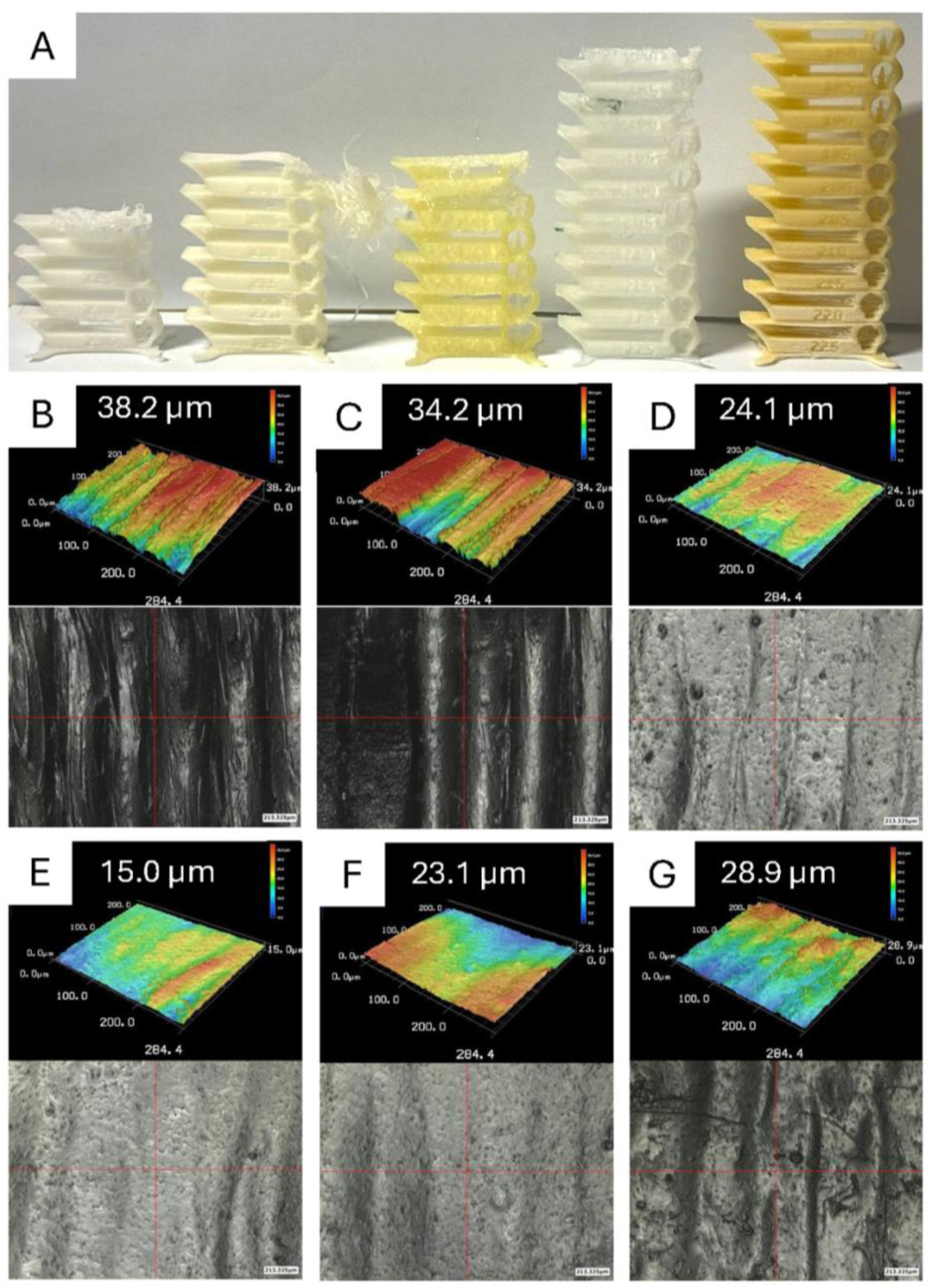
Screening of soluble filaments and surface-smoothing strategies for 3D-printed sacrificial cores. (**A**) Exemplars of the five candidate soluble filaments (left to right): Xioneer VXL90 (lye-soluble copolymer), BASF Ultrafuse BVOH, Formfutura AquaSolve PVA, MatterHackers PRO Series PVA, and eSUN PVA. (**B - G**) Scanning laser microscopy of eSUN PVA test coupons printed in spiral-vase mode (layer height 0.05 mm; extrusion width 0.6 mm) to quantify surface roughness. Treatments: (**B**) no surface modification; (**C**) 1200-grit sandpaper; (**D**) wet laboratory wipe (KimWipe); (**E**) 30 s dip in static room-temperature water; (**F**) 1 min dip in static room-temperature water; (**G**) 3 min dip in static room-temperature water. The numeric labels above each panel indicate the measured mean roughness (µm) for the imaged field. **B-G** Scale Bars = 213.325µm.

**Supplementary Figure 2:**
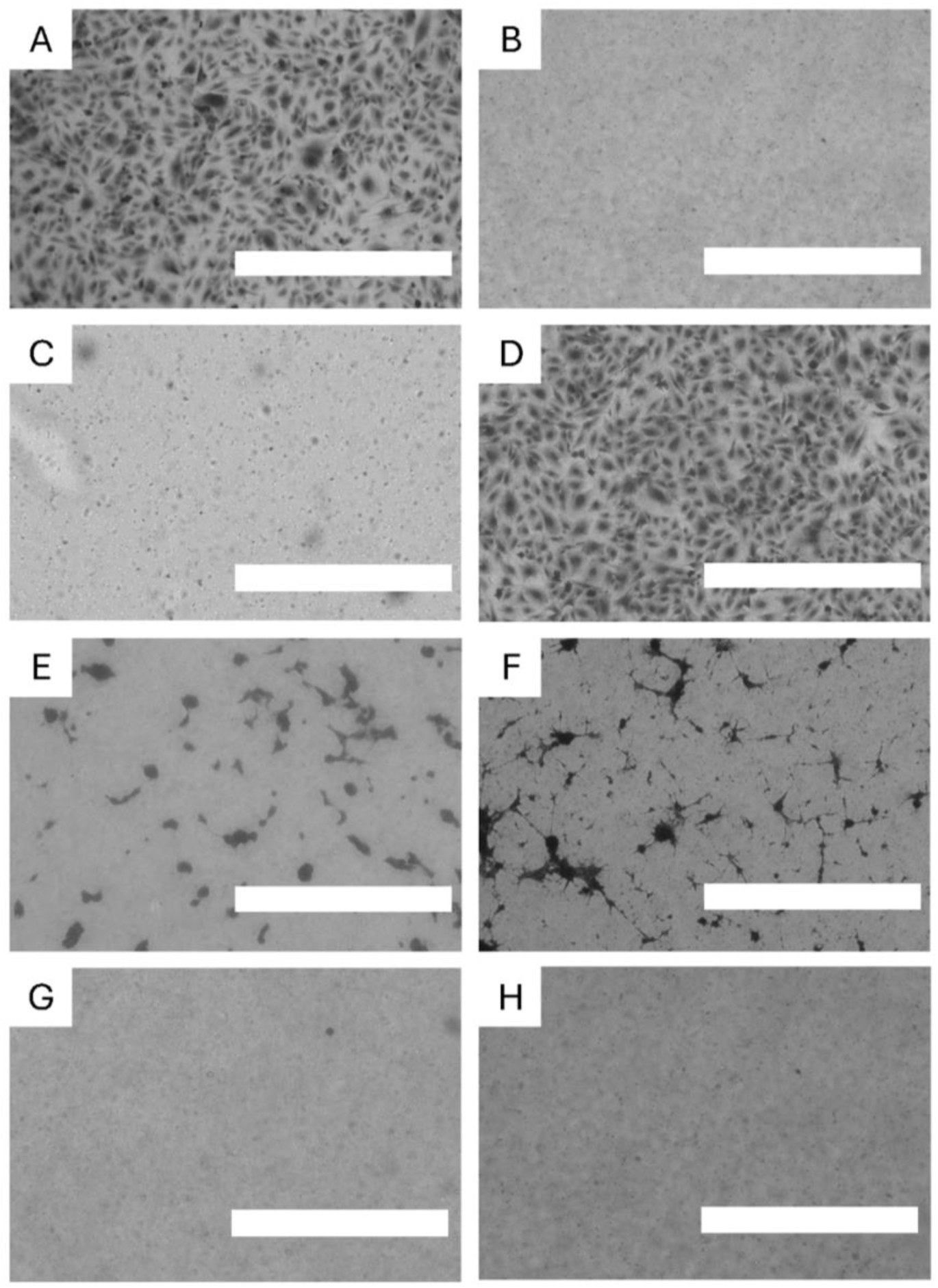
Optimizing HMEC-1 attachment on PDMS. Representative brightfield micrographs of HMEC-1 endothelial cells stained with Crystal Violet after 24 h of growth on different modified surfaces: (**A**) standard tissue-culture–treated plate (positive control); (**B**) unmodified PDMS; (**C**) PDMS + 1% gelatin; (**D**) PDMS + fibronectin (10 μg/mL); (**E**) PDMS + fibronectin (10 μg/mL) with 1% Pluronic and 1% PEG-PDMS; (**F**) PDMS subjected to plasma treatment then coated with fibronectin (10 μg/mL); (**G**) PDMS + fibronectin (10 μg/mL) with 1% Pluronic; (**H**) PDMS + fibronectin (10 μg/mL) with 1% PEG-PDMS. Scale bars = 1 mm.

**Supplementary Figure 3:**
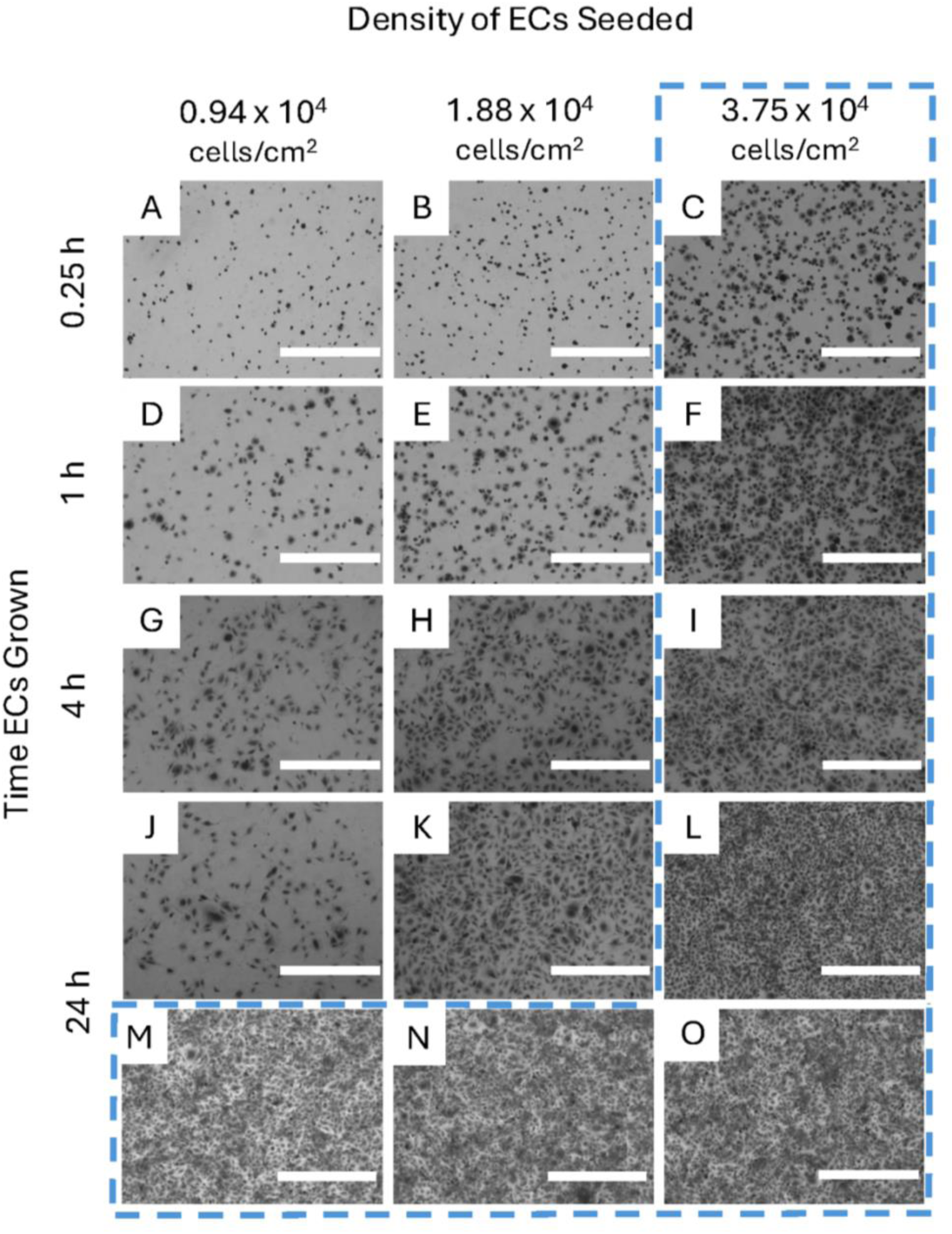
Optimizing HMEC-1 seeding density and fibronectin incubation for rapid monolayer formation. Representative brightfield micrographs of HMEC-1 endothelial cells grown on PDMS pre-coated with fibronectin, fixed and stained with crystal violet. Cells were grown for 0.25 h (**A–C**), 1 h (**D–F**), 4 h (**G–I**), 24 h (**J–0**). At 24 h, surfaces additionally pre-incubated with fibronectin for varying durations showed similar outcomes at the highest seeding density: M (1 h coat), N (2 h coat), O (4 h coat), all 3.75×10⁴ cells/cm². Increasing seeding density accelerated coverage; the highest density (C, F, I, L, M–O) produced near-confluent monolayers by 24 h. The protocol derived from these data: fibronectin coat 1–24 h, seed 3.75×10⁴ cells/cm², and culture ≥24 h to obtain a robust endothelial monolayer. Scale bars, 1 mm.

**Supplementary Figure 4:**
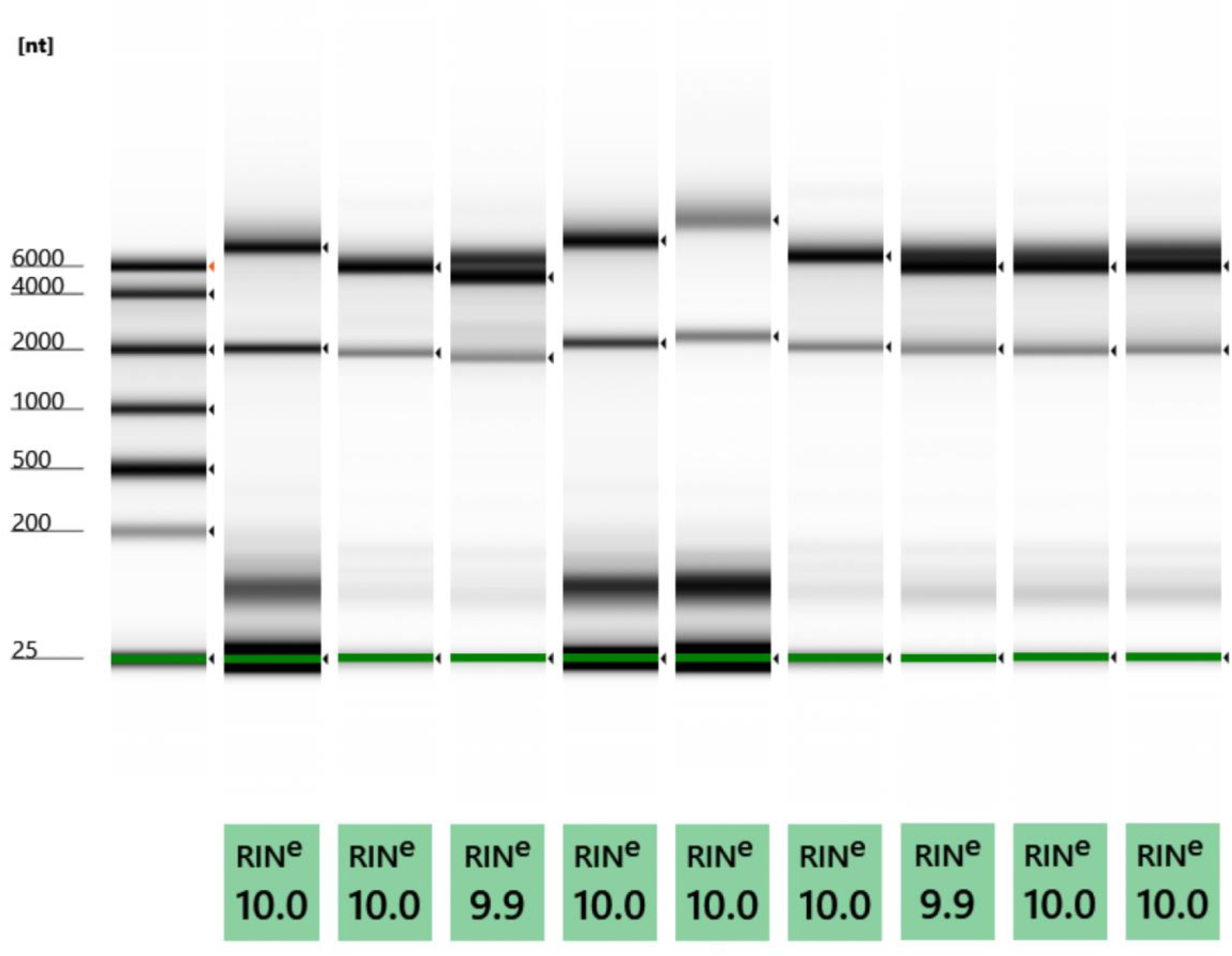
RNA integrity across static, continuous, and pulsatile flow conditions. Capillary electrophoresis showing the RNA ladder (lane 1; sizes in nucleotides at left), followed by triplicate HMEC-1 samples from static culture (lanes 2–4), continuous flow (lanes 5–7), and pulsatile flow (lanes 8–10). All samples display sharp 18S/28S rRNA bands with minimal low-molecular-weight smear, indicating excellent integrity. Estimated RNA Integrity numbers (RINe) were 9.9–10.0 for all biological replicates.

**Supplementary Figure 5:**
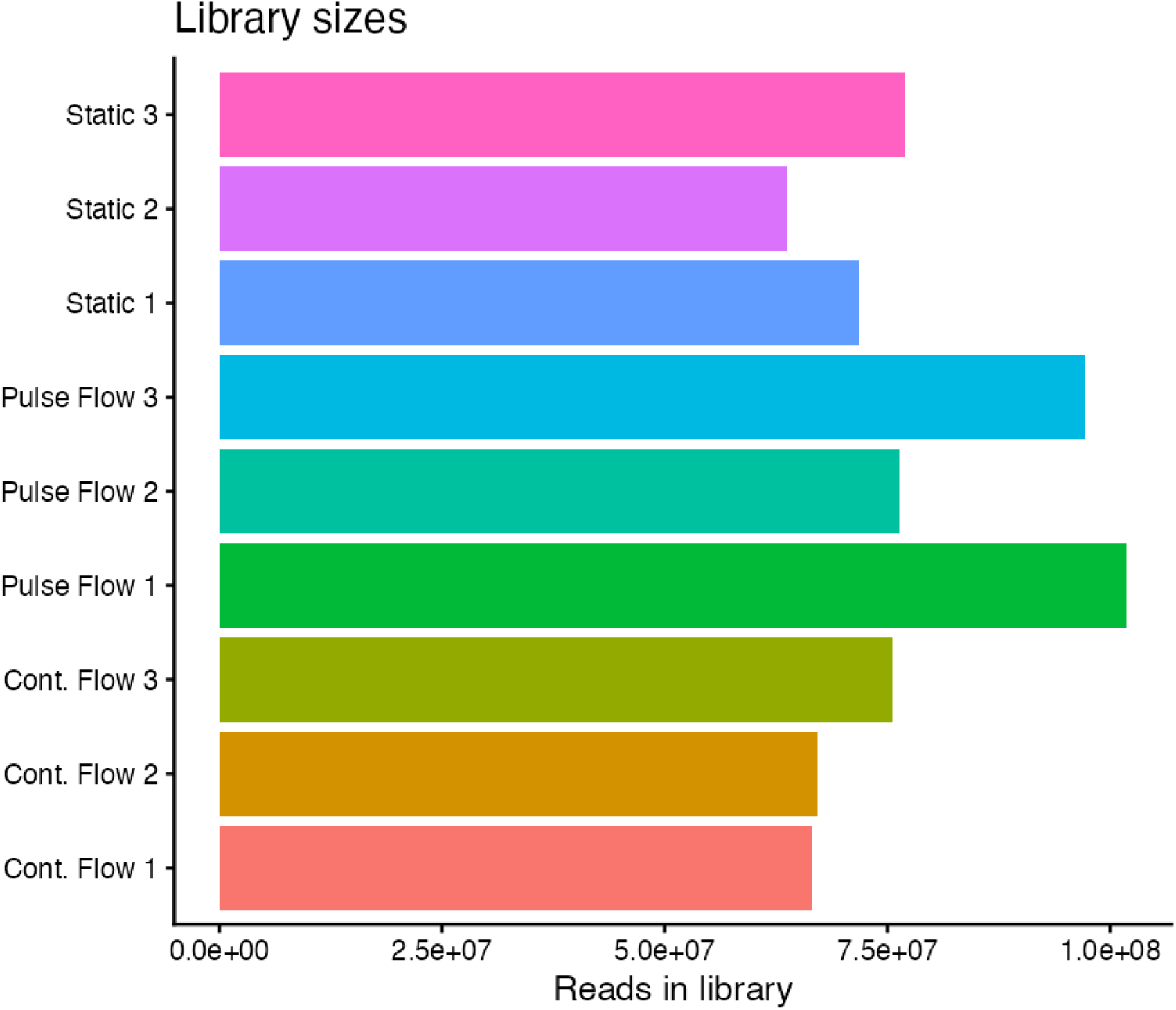
RNA-seq library sizes across samples. Horizontal bar chart showing total reads per library for all nine samples: Static 1–3, Continuous Flow (Cont. Flow) 1–3, and Pulsatile Flow (Pulse Flow) 1–3. Libraries range from ∼6.6×10^7 to ∼1.0×10^8 reads, with broadly comparable depths across conditions. Similar sequencing yields indicate balanced coverage suitable for downstream normalization and differential expression analyses.

**Supplementary Figure 6:**
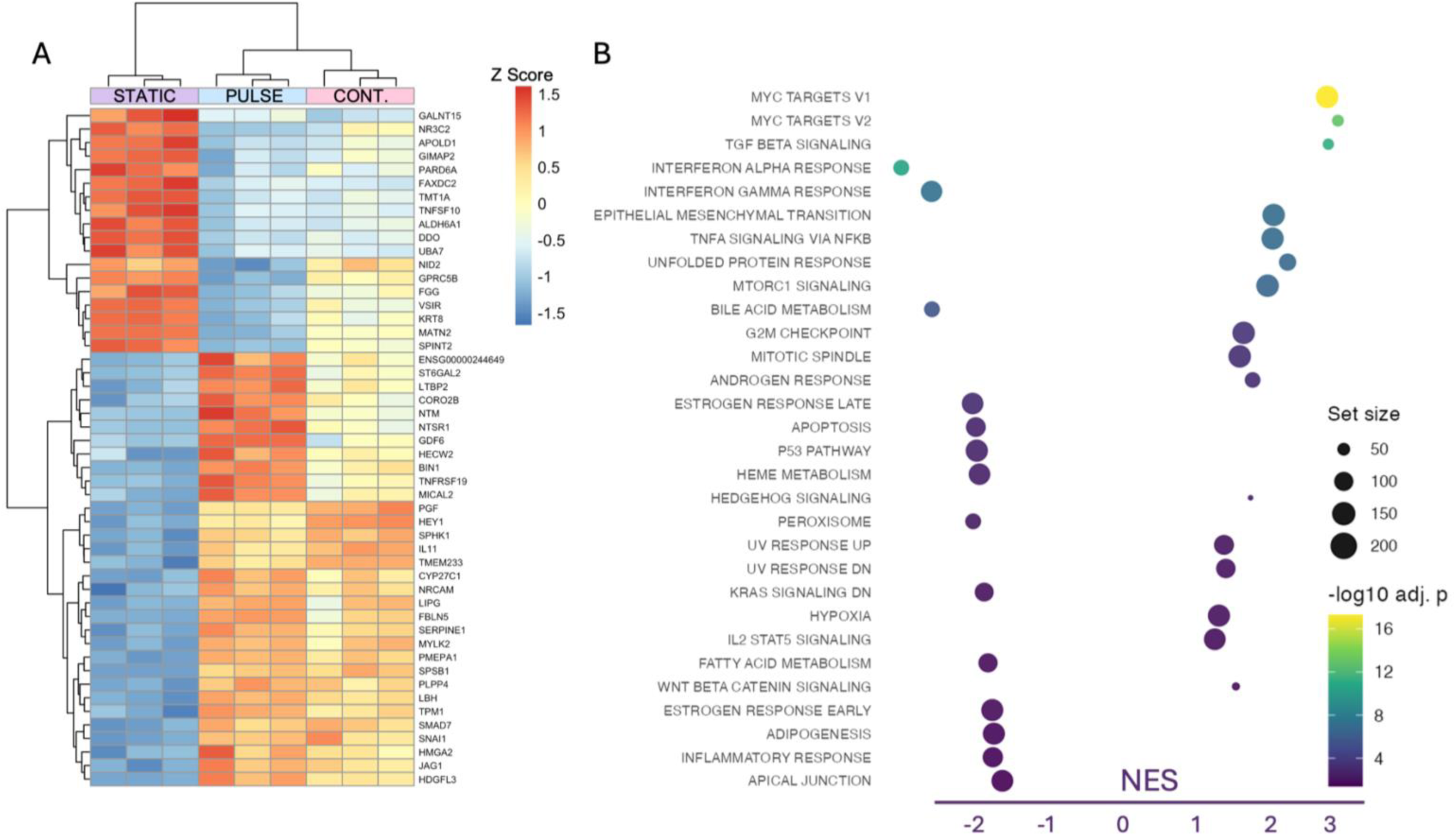
Heat map of the top 50 differentially expressed genes in pulsatile flow vs. static conditions comparison across all samples (columns). **(A)** Values are variance-stabilized counts (DESeq2) and z-scored by gene. Rows and columns were hierarchically clustered using 1–Pearson correlation distance and complete linkage. **(B)** Normalised enrichment score (NES) for Pulsatile vs static flow (positive = enriched in pulsatile flow; negative = enriched in static culture).

**Supplementary Figure 7:**
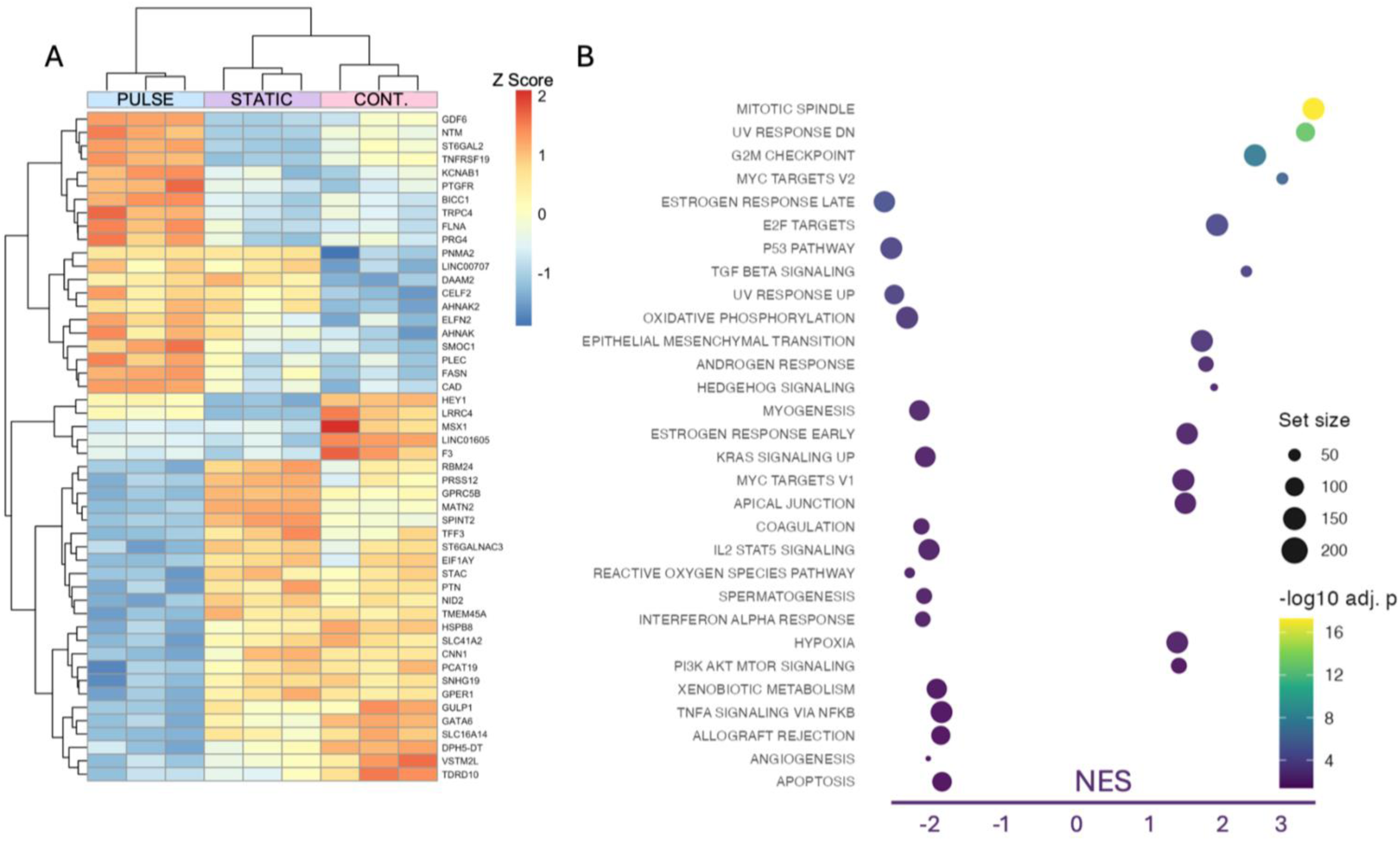
Heat map of the top 50 differentially expressed genes in pulsatile flow vs. continuous flow comparison across all samples (columns). **(A)** Values are variance-stabilized counts (DESeq2) and z-scored by gene. Rows and columns were hierarchically clustered using 1–Pearson correlation distance and complete linkage. **(B)** Normalised enrichment score (NES) for Pulsatile vs Continuous flow (positive = enriched in pulsatile flow; negative = enriched in continuous flow).

